# Genomic insights into the phylogeny and biomass-degrading enzymes of rumen ciliates

**DOI:** 10.1101/2022.01.05.474053

**Authors:** Zongjun Li, Xiangnan Wang, Yu Zhang, Zhongtang Yu, Tingting Zhang, Xuelei Dai, Xiangyu Pan, Ruoxi Jing, Yueyang Yan, Yangfan Liu, Shan Gao, Fei Li, Youqin Huang, Jian Tian, Junhu Yao, XvPeng Xing, Tao Shi, Jifeng Ning, Bin Yao, Huoqing Huang, Yu Jiang

**Affiliations:** Center for Ruminant Genetics and Evolution, College of Animal Science and Technology, Northwest A&F University, Yangling 712100, China; Department of Animal Sciences, The Ohio State University, Columbus, OH 43210, USA; Key Laboratory of Agricultural Animal Genetics, Breeding, and Reproduction of Ministry of Education, College of Animal Science and Technology, Huazhong Agricultural University, Wuhan, 430070, China; State Key Laboratory of Grassland Agro-ecosystems, College of Pastoral Agriculture Science and Technology, Lanzhou University, Lanzhou, 730020, China; State Key Laboratory of Animal Nutrition, Institute of Animal Sciences, Chinese Academy of Agricultural Sciences, Beijing 100193, China; Key Laboratory of Animal Biotechnology of the Ministry of Agriculture, College of Veterinary Medicine, Northwest A&F University, Yangling, 712100, China; College of Information Engineering, Northwest A&F University, Yangling, 712100, China

## Abstract

Understanding the biodiversity and genetics of the gut microbiome has important implications for host physiology. One underexplored and elusive group is ciliated protozoa, which play crucial roles in regulating gut microbial interactions. Integrating single-cell sequencing and an assembly-and-identification pipeline, we acquired 52 high-quality ciliate genomes of 22 rumen morphospecies for all major abundant clades. With these genomes, we firstly resolved the taxonomic and phylogenetic framework that reclassified them into 19 species spanning 13 genera and reassigned the genus Dasytricha from Isotrichidae to a new family Dasytrichidae. Via extensive horizontal gene transfer and gene family expansion, rumen ciliates possess a broad array of enzymes to synergistically degrade plant and microbial carbohydrates. In particular, ∼80% of the degrading enzymes in Diplodiniinae and Ophryoscolecinae act on plant cell wall, and the high activities of their cellulase, xylanase and lysozyme reflect the potential of ciliate enzymes for biomass-conversion. Additionally, the new ciliate dataset greatly facilitated the rumen metagenomic analyses by allowing ∼12% of reads to be classified.

## Introduction

The genomes of hundreds of thousands of gut prokaryotes have been sequenced, which have greatly advanced our understanding of their taxonomic and functional diversity and important roles in host health and nutrition^1–3^. Trichostome ciliates (of the phylum Ciliophora, class Litostomatea) are ubiquitous in the gut microbiome of vertebrates^4,5^, and they serve crucial roles in modulating the gut microbiome through predation, competition, and symbiosis^6^. However, genomic research on gut ciliates has lagged^1^, and by far only *Entodinium caudatum*, one of the most common species in the rumen of ruminants, has been subjected to genomic study^7^. The lack of genomic information on the ciliate has become an obstacle for holistic studies of the gut microbiome, rumen microbiome in particular, as ciliates can account for up to 50% of the total rumen microbial biomass and approximately 200 morphospecies have been described in the rumen^6^.

Despite their prominence in the rumen ecosystem and extensive previous research over one century^8^, much remains to be learned about rumen ciliates^9,10^. Based on morphological features, rumen ciliates are primarily classified into two clades: family Isotrichidae in the order Vestibuliferida (with cilia covering the entire cell surface) and family Ophryoscolecidae in the order Entodiniomorphida (with localized zones of cilia including one or two adoral cilia zone)^4,6^. It has been recognized, however, taxonomic identification and classification of ciliates based on morphological features are challenging^6,11^, as the same species can exhibit a few different morphotypes^12^. The 18S rRNA gene of ciliates has been used to aid species identification^13^, but it has several limitations in proposing a new taxonomic framework, including topological discrepancies even at the subfamily level^5,14^ and too conserved to uncover cryptic species^15^.

It has also been challenging to determine or investigate the actual metabolism of rumen ciliates because of the lack of and inability to establish axenic cultures. As a result, our knowledge of rumen ciliate metabolism can only be inferred from studies using monocultures (that contain bacteria and other microbes) or defaunation (complete removal of rumen protozoa chemically or through physical isolation), both of which introduce confounding effects of other microbes and residual effects^10,16^. Recent genomic and transcriptomic studies provided new insights into the metabolism and physiology of several species of rumen ciliates^7,17^. However, the functional diversification and evolutionary mechanisms underlying most rumen ciliates linkages remain undetermined. Additionally, the unique genome structures (e.g., nuclear dimorphism, chromosomal fragmentation) make ciliates valuable model organisms to understand many important fundamental molecular and cellular processes and features, such as catalytic RNA, telomerase, and histone acetylation^18,19^. However, most knowledge comes from a few free-living aerobic ciliate species within the classes Oligohymenophorea and Spirotrichea^20,21^, while rumen ciliates, which are anaerobic and symbiotic, are exclusively found in the class Litostomatea.

In this study we *de novo* assembled and analyzed 52 high-quality single-ciliate-cell amplified genomes (SAGs) from 22 rumen morphospecies, which were reclassified into 19 species spanning 13 genera and three families in the newly genome-based taxonomy. One assembly-and-identification pipeline was developed to help improve the quality and accuracy of genome assembly of ciliates. This unparalleled genome catalog of rumen ciliates greatly facilitated and/or enabled better understanding and determination of their taxonomy, phylogeny, metabolism, and ecology.

## Results

### Recovering single-ciliate-cell amplified genomes

We sequenced the DNA amplified from 69 individual single ciliate cells isolated from fresh rumen fluid sampled from 15 cattle and goats (**Supplementary Table 1**). They represented 22 ciliate morphospecies (**Fig. 1** and **Supplementary Fig. 1-3**) belonging to 11 prevalent and abundant morphogenera, which represent ∼97% of the genus-level abundance across our collected samples and published studies (**Supplementary Table 2**). We also sequenced the amplified transcriptome^22^ of 24 of the isolated single-ciliate-cells and 14 rumen metatranscriptomes for *de novo* gene prediction (**Supplementary Fig. 4**).

**Fig. 1.**
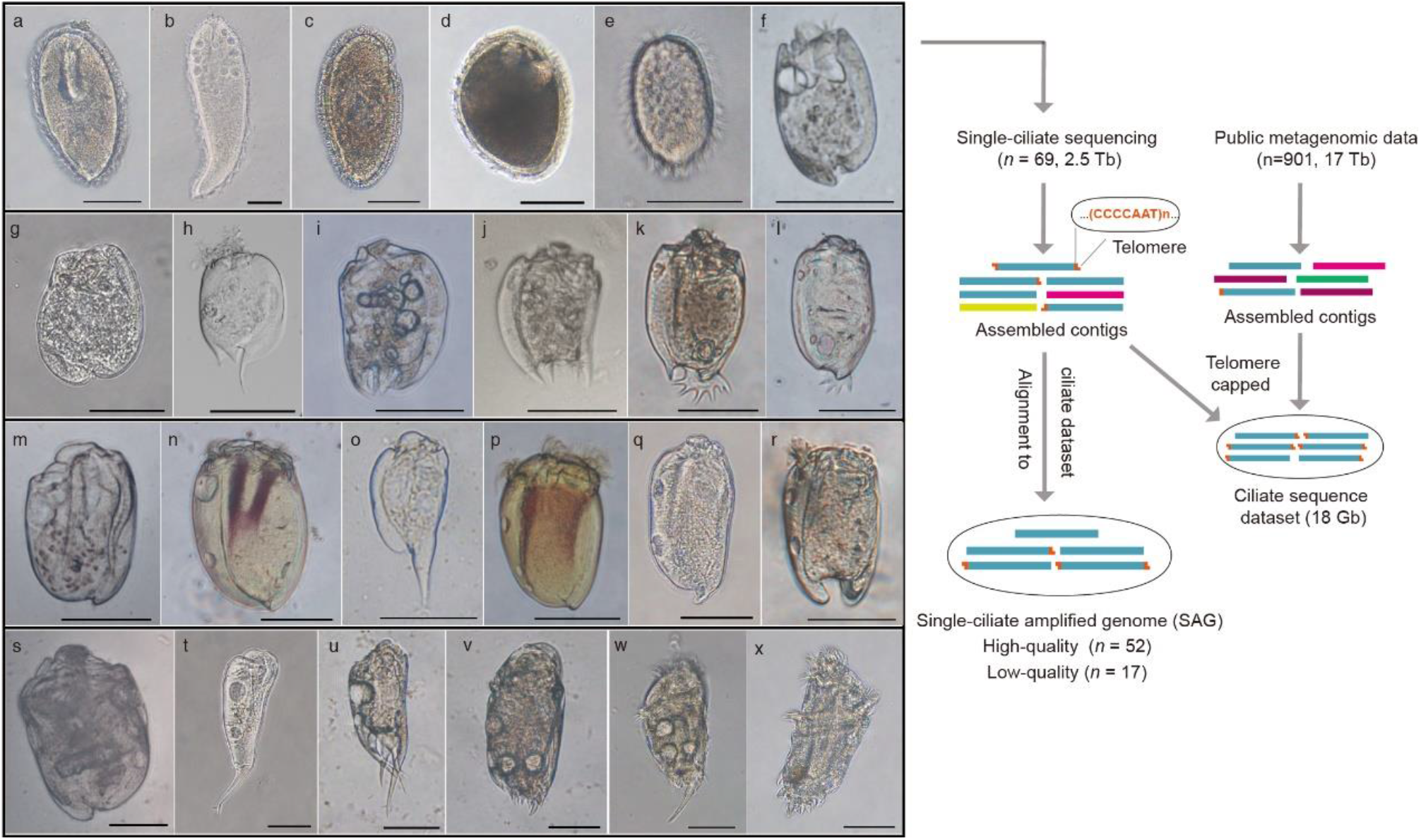
Light micrographs of 22 rumen ciliate morphospecies and two cryptic species examined in this study, and a brief pipeline for recovering single-ciliate genomes. **a**, *Iso. prostoma*. **c**, *Iso. intestinalis*. **e**, *Das. ruminantium*. **f**, *Ent. longinucleatum*. **g**, *Ent. bursa*. **h**, *Ent. caudatum*. **i**, *Dip. anisacanthum*. **j**, *Dip. dentatum*. **k**, *Dip. flabellum monospinatum*. **l**, *Dip. flabellum aspinatum*. **m**, *Eno. triloricatum*. **n**, *Met. minomm*. **o**, *Ere. rostratum*. **p**, *Ost. gracile*. **q**, *Ost. venustum*. **r**, *Ost. mammosum*. **s**, *Pol. multivesiculatum*. **t**, *Epi. caudatum*. **u**, *Epi. cattanei*. **v**, *Oph. bicinctus*. **w**, *Oph. caudatus*. **x**, *Oph. purkynjei. Iso. sp. YL-2021a* (**b)** and *Iso. sp. YL-2021b* (**d**) were two cryptic species of Isotrichidae. Micrographs **n** and **p** show stained skeletal plates. Scale bars, 50 µm (a-x). More details of the rumen ciliates are shown in **Supplementary Fig. 1**.

Because rumen ciliate cells carry other microbes as endosymbionts or engulfed microbes (primarily bacteria) and because the DNA of the latter can also be amplified and sequenced, we developed an assembly-and-identification pipeline to recover and identify ciliate macronuclear sequences (**Fig. 1** and **Supplementary Fig. 4**). Briefly, using contigs assembled from the 2.5 Tb sequences from the single cells and the ∼17 Tb publicly available rumen metagenome (**Supplementary Table 3**), we constructed an 18 Gb assembled dataset of contigs of rumen ciliate macronuclear, all of which were capped on both or one end with telomeric repeats ((CCCCAAT)n). This dataset was searched for the contigs that did not have any telomeres in the single-ciliate assembled contigs (see **Methods**). Additionally, we performed hybrid assembly and deep sequencing to improve the assembly quality (**Supplementary Fig. 5**). Of the 69 SAGs we assembled (**Supplementary Table 4**), 52 had ≥80% of the 171 Alveolata conserved marker genes^23^ and were considered “high-quality”, while 17 had <80% of the Alveolata conserved marker genes. The 52 high-quality SAGs were used for further analysis. Among the 52 SAGs, the telomere-capped contigs ranged from 14% to 92% (mean 61%) of the total contigs (**Supplementary Table 4**). One of the SAGs of *Oph. caudatus* (referred to as SAGT3) had the highest number (26,497 to be exact) of complete chromosomes (contigs capped with telomere at both ends), plus another 33,578 contigs that were capped with telomere at only one end. The chromosome number of *Oph. caudatus* is at least 43,286, which is much more than the known large number of chromosomes in ciliates, *Oxytricha trifallax* (16,000) and *Halteria grandinella* (23,000)^20,24^.

### Deriving a robust genome-based taxonomy and phylogeny

A robust taxonomy is an organizing principle of biology to communicate scientific results and describe biodiversity^25,26^. Using the 52 SAGs, we established a robust genome-based taxonomic framework of rumen ciliates that overcomes the uncertainty inherent of the morphology-based taxonomy.

Whole-genome average nucleotide identity (ANI) has emerged as a widely accepted method for circumscribing prokaryotic species, with 95% ANI as a species boundary^26,27^. We found a clear ANI discontinuity among the SAGs: >97% or <92% (**Fig. 2a-b**) and considered ANI >97% and <92% for intra-species and inter-species cutoffs, respectively. Based on the above ANI cutoff values, we found four pairs of synonymic species in Ophryoscolecidae (**Fig. 2a** and **Supplementary Fig. 6**). This is exemplified by *Oph. caudatus, Oph. purkynjei* and *Oph. bicinctus* (**Fig. 1v-x** and **Supplementary Fig. 3**) being classified as a single species, although *Oph. caudatus* was isolated from sheep and goats with a long main caudal spine and three circles of secondary caudal spines, whereas *Oph. purkynjei* and *Oph. bicinctus* were isolated from cattle with a short main caudal spine and two circles of secondary caudal spines, respectively^6^. Similarly, *Dip. anisacanthum* and *Dip. dentaum, Dip. flabellum aspinatum* and *Dip. flabellum monospinatum, Ost*.*gracile* and *Ost. venustum* were synonymy, respectively (**Fig. 2a** and **Supplementary Fig. 1**). These suggest that the shape of the main caudal spine, the number of secondary caudal spines, and habitat host are not reliably taxonomic worth for Ophryoscolecidae. On the other hand, despite having morphological similarity, two new cryptic species of Isotrichidae were identified (ANI < 92%, **Fig. 2a**): tentatively named as *Iso. sp. YL-2021a* and *Iso. sp. YL-2021a*. The former was a cryptic species (**Fig. 1b**) of *Iso. prostoma* and it had a longer body (278 µm) than *Iso. prostoma*, while the latter was a cryptic species (**Fig. 1d**) of *Iso. intestinalis* and it had a lower ratio of body length to width (1.3 on average) but longer chromosomes (22.5 kb on average) than *Iso. intestinalis*. Additionally, the 18S sequence identity between synonymic morphospecies was all >99% while that among the four Isotrichidae species was all <98% (**Supplementary Fig. 6**), which also supports the taxonomic revision at the species level. Accordingly, the 22 collected morphospecies were reclassified to 19 species.

**Fig. 2.**
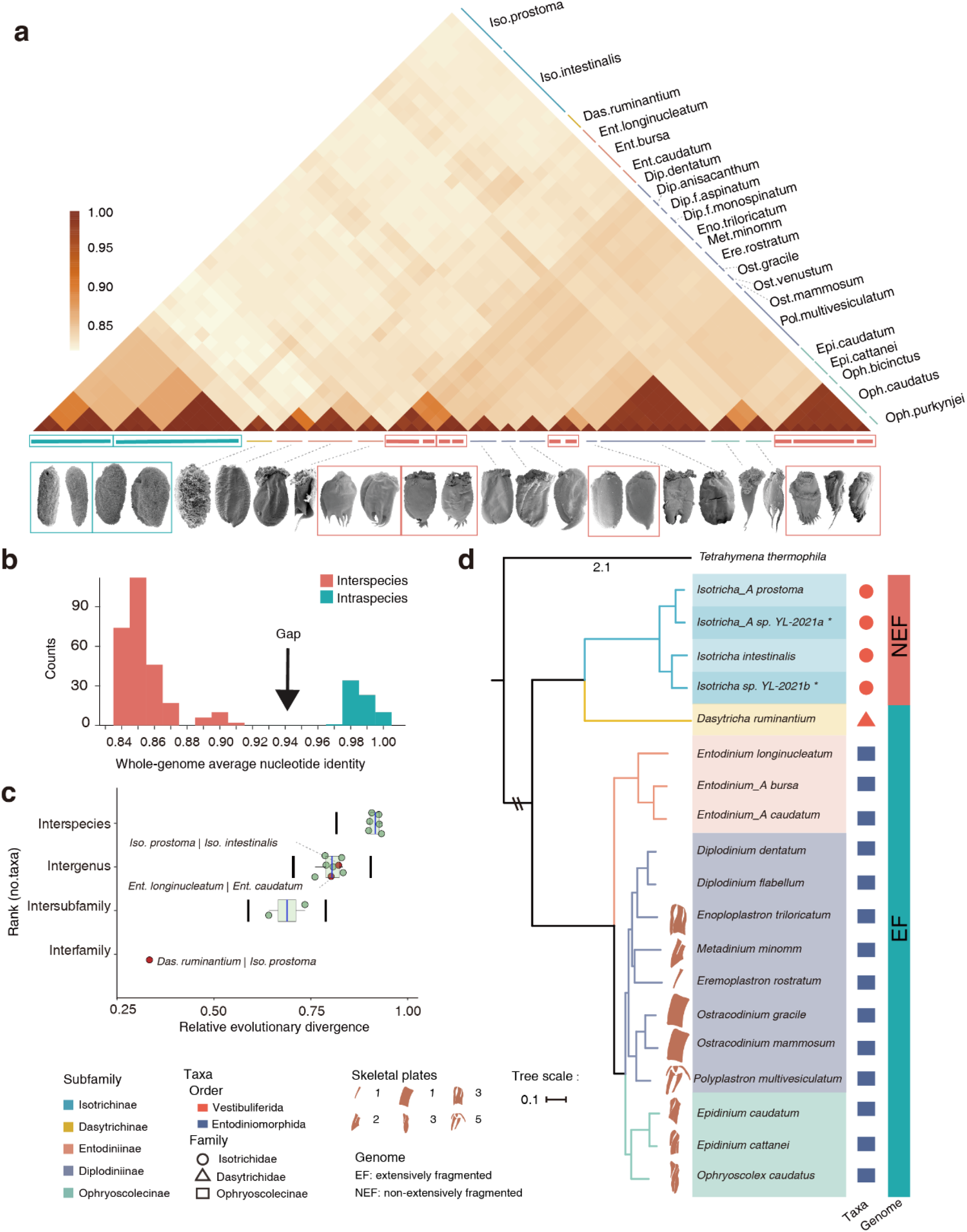
Genome-based taxonomy and phylogeny of rumen ciliates. **a**, A heatmap showing the ANI among the 52 SAGs and the scanning electron microscopy images of 22 corresponding morphospecies and two cryptic species. **b**, ANI distributions. **c**, Rank normalization through relative evolutionary divergence (RED) of taxa. The red dot represents rank reassignment. The RED interval for each rank is shown by two vertical black lines, median±0.1. **d**, The maximum likelihood phylogenetic tree of 19 rumen ciliate species and *T. thermophila* (as the outgroup) based on 113 concatenated single-copy proteins. All nodes have 100% bootstrap support. The black asterisks indicate the two cryptic species identified in this study.

The current taxonomic ranks for rumen ciliates at or above the genus level also lack molecular evidence. For example, the 18S rRNA gene phylogeny placed the subfamilies Entodiniinae and Ophryoscolecinae as two branching lineages within the subfamily Diplodiniinae^5,14^. To resolve the phylogenetic challenges, we firstly constructed a maximum likelihood (ML) tree using 113 concatenated single-copy proteins with *Tetrahymena thermophila* as an outgroup (**Fig. 2d**). We also constructed a species tree using OrthoFinder^28^ under the coalescent model (**Supplementary Fig. 7**). The two trees were consistent in topology and had 100% bootstrap support for each node. The results support Entodiniinae representing the oldest branch in Ophryoscolecidae and Ophryoscolecinae as a sister to Diplodiniinae, which is consistent with their morphological inference^29^. The concatenated protein-based phylogeny served as the basis for the taxonomic rank normalization at and above genus using relative evolutionary divergence (RED)^25^. Comparing the RED value with the RED intervals of taxonomic ranks (median RED ± 0.1), the genus Dasytricha was reassigned from Isotrichidae to a new family Dasytrichidae, which was treated as a synonymy of Isotrichidae in the past^4^. Additionally, *Ent. bursa* and *Ent. caudatum* were reassigned to a new genus Entodinium_A, and *Iso. prostoma* and *Iso. sp. YL-2021a* were reassigned to a new genus Isotricha_A (**Fig. 2c-d**). For consistency with the family Ophryoscolecidae, subfamilies Isotrichinae and Dasytrichinae were created for the families Isotrichidae and Dasytrichidae, respectively. Ultimately, the 19 identified ciliate species were assigned to two orders, three families, five subfamilies, and 13 genera (**Fig. 2d**). The genome-based taxonomy supports the taxonomic worth of the size and number of skeletal plates, which is a newly evolved organelle for storing amylopectin in the clades of Ophryoscolecinae and Diplodiniinae^6^ and useful in differentiating genera. However, the phylogenetic relationships among the genera within Diplodiniinae did not reflect the number of skeletal plates, as evidenced by Diplodinium without skeletal plate but being more closely related to Enoploplastron with three skeletal plates than Eremoplastron with only one skeletal plate. This suggests a single origin of the skeletal plates, which was probably lost in Diplodinium.

Invasion into the rumen was one of the most important events in the evolutionary history of Trichostome ciliates^5^. As shown above, not only rumen Ophryoscolecidae, as previously suggested^5^, but also Isotrichidae have radiated in speciation. Molecular clock analyses showed that explosive radiation of both Ophryoscolecidae and Isotrichidae occurred about 5 - 35 million years ago (**Fig. 3e** and **Supplementary Fig. 8**), which is consistent with the duration of the rapid radiation of ruminants^30^, suggesting concurrent co-evolution and co-speciation between ruminants and rumen ciliates.

**Fig. 3.**
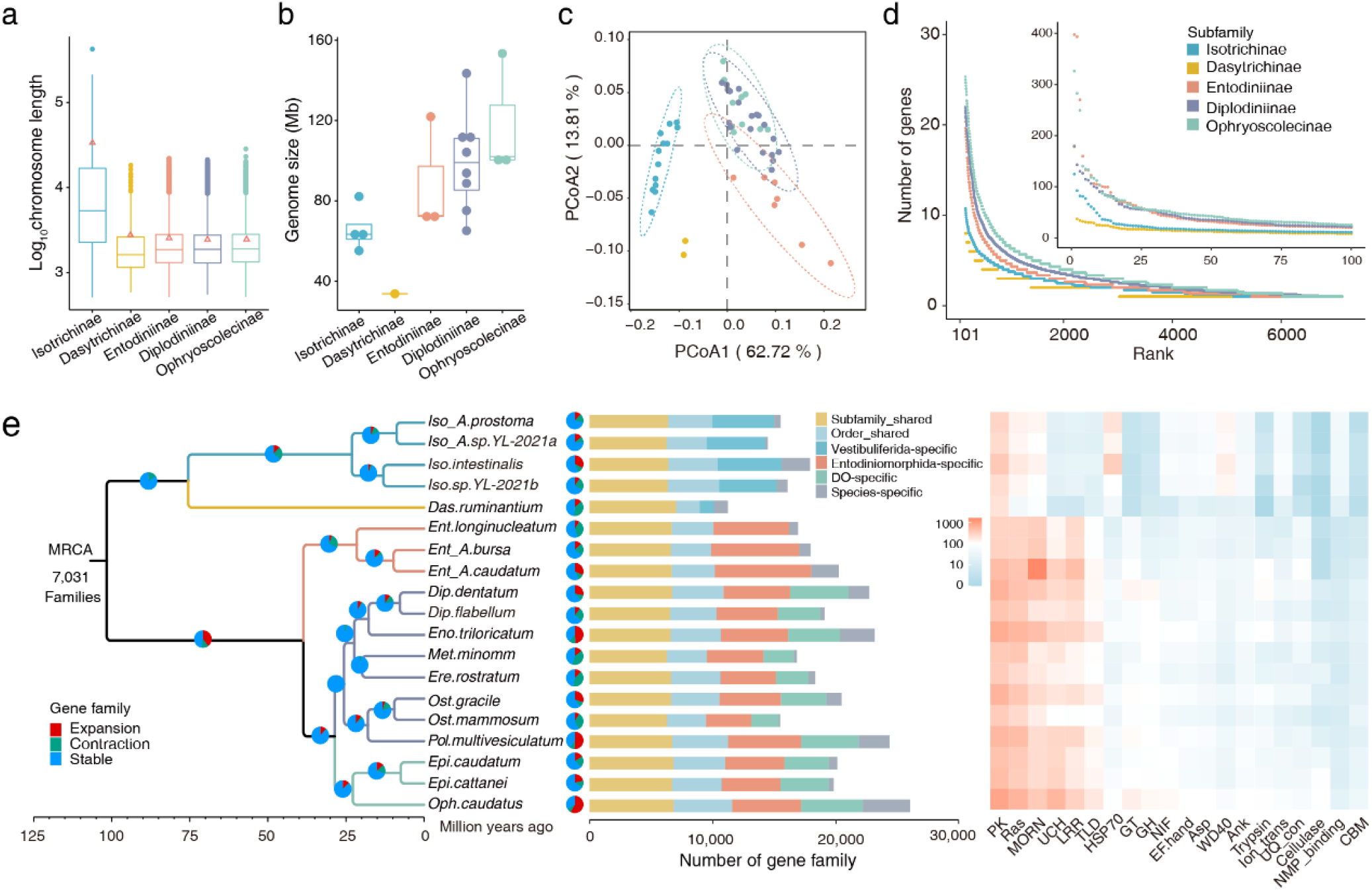
Genome characteristics of rumen ciliates. **a** Chromosome length (the red triangles indicate N50) and **b** genome size distributions across five subfamilies. **c** Functional divergence based on the clans of Pfam (P-value of Adonis < 0.001). **d** Rank curves showing gene number divergence in the subfamilies-shared gene families. **e** Gene family divergence: From left to right: a time phylogenetic tree of 19 rumen ciliate species with gene family expansions and contractions; comparison of the gene family repertoires among the 19 ciliate species; a heat map showing the top 10 categories of domains found in order-specific gene families of each species.

### Genome architecture and gene family diversity

Explosive speciation can result from a series of genomic changes and innovations, and the genome catalog of the rumen ciliates provides new insights into their early diversification.

The genomic architecture features of Isotrichidae were obviously divergent from those of Dasytrichidae and Ophryoscolecidae (**Supplementary Table 4**), despite Dasytrichidae and Isotrichidae being in the same order. Notably, Isotrichidae had a lower GC content and longer gene-coding regions and gene introns than Dasytrichidae and Ophryoscolecidae (**Supplementary Fig. 9-10**). Most interestingly, the two classic chromosome architectures of ciliates macronuclear genomes^31^ existed in a single subclass (**Fig. 3a**): the genomes of Ophryoscolecidae and Dasytrichidae were extensively fragmented (EF) with gene-sized nanochromosomes having a mean length of 2.3 kb, while the genomes of Isotrichidae were non-extensively fragmented (NEF) minichromosomes with a mean length of 14.3 kb, mostly encoding several genes each. The genomes of its sister-subclasses Haptoria and sister-classes Spirotrichea were both EF^32^, therefore, we inferred that the genome of the common ancestor of rumen ciliates was EF, and the NEF genomes of Isotrichidae was an independent origin according to the parsimony principle of evolution^33^. However, similar to EF genomes, the chromosome copy numbers in the NEF genomes of Isotrichidae were nonuniform (**Supplementary Fig. 11**), while those in the NEF genomes of the classes Heterotrichea and Oligohymenpphores were approximately equal^32^.

Using OrthoFinder^28^, we annotated 64,189 gene families from the 52 SAGs (**Fig. 3e** and **Supplementary Fig. 10**), of which, 40,898 were shared by at least two species, and 23,291 were identified as species-specific (ranging from 110 in *Ost. mammosum* to 3,818 in *Oph. caudatus*). Of the species-shared families, only 14,871 (36.4%) were found in both Vestibuliferida and Entodiniomorphida, while the remaining gene families were Vestibuliferida-specific (6,091) or Entodiniomorphida-specific (19,936). Of the Entodiniomorphida-specific families, 10,043 arose at the clades of Diplodiniinae and Ophryoscolecinae. This suggests that extensive lineage-specific gene families arose in the early and during the diversification of rumen ciliates into modern orders. Additionally, compared with the common ancestor of rumen ciliates, more subfamily-shared gene families expanded in Entodiniomorphida (2,784/7,031) than in Vestibuliferida (53/7,031) (**Fig. 3e**). As a result, species in Entodiniomorphida, in particular in the clade of Diplodiniinae and Ophryoscolecinae, had larger and more diverse gene families than the species in Vestibuliferida (**Fig. 3d**), which might have further induced the functional divergence across orders and subfamilies (**Fig. 3c**). Comparing the differential investment in KEGG pathways and Gene Ontology terms across those clades (**Supplementary Fig. 12-13**), we found that the members of the Isotrichidae possessed a higher proportion of genes related to organismal systems and environmental adaptation (e.g., endocrine, immune, detoxification, and antioxidant activity), while those of Ophryoscolecidae were primarily related to biomass (e.g., carbohydrates, lipids, and xenobiotics) degradation and metabolism. Indeed, compared the domains of order-specific genes between Isotrichidae and Ophryoscolecidae, Isotrichidae had a relative higher predominance of HSP70 and WD40, which are related to stress resistance^34^ and biogenesis of motile cilia^35^, respectively, while Ophryoscolecidae had a higher relative predominance of CAZymes-related domains (GT, GH, CBM and cellulase) (**Fig. 3e**).

### Rumen ciliates possess a broad array of CAZymes

The gut microbes are of huge industrial interest due to their ability to release fermentable sugars from lignocellulose^36–38^. Here, we found that the rumen ciliates were also a rich source of biomass-degrading enzymes. A total of 33,693 non-redundant (< 99% identity) CAZymes were predicted in the 52 SAGs (**Fig. 4a**), and only 357 had a highly similar match (≥95% identity) in any of the public protein databases (nr, env_nr, m5nr, UniProt TrEMBL) and the rumen bacterial and fungal protein datasets^1,2,39,40^, indicating that 99% of the CAZymes can be considered new. In particular, the species of Diplodiniinae and Ophryoscolecinae encoded as many CAZyme as gut fungi, the latter of which possesses the largest number of CAZyme genes yet known in nature^37^. And the species of Diplodiniinae and Ophryoscolecinae encoded more CAZyme genes than Entodiniinae (by two-fold) and Isotrichidae and Dasytrichidae (by five-fold) (**Fig. 4b**) with clear profile-divergences (**Fig. 4c**), largely attributed to the extensive gene family arising and expansion.

**Fig. 4.**
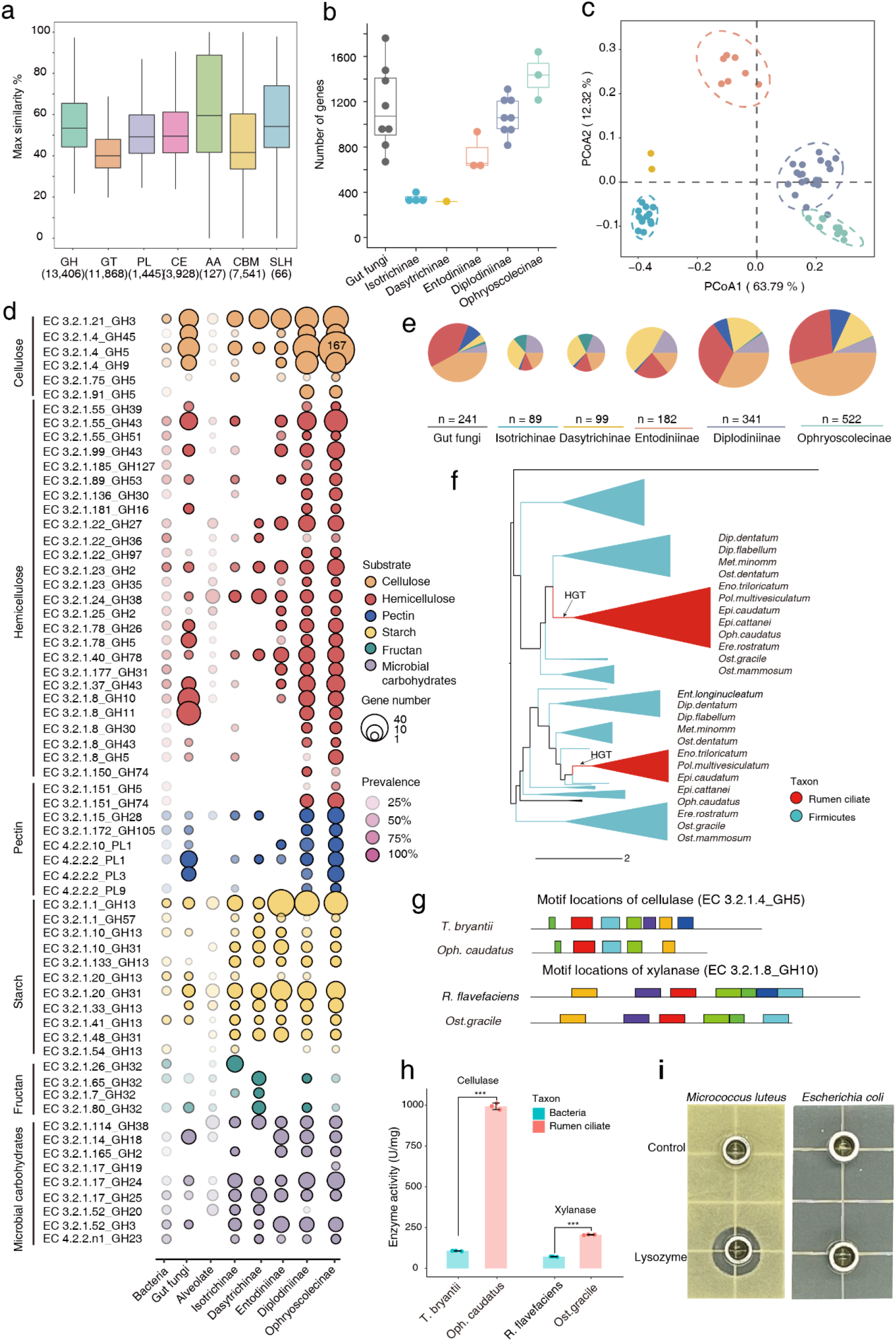
CAZyme profiles of rumen ciliates. **a**, Max similarities (amino acid sequences) of the CAZymes identified in the rumen ciliates compared to the public databases for seven classes of CAZymes. GH, glycoside hydrolase; GT, glycosyl transferase; PL, polysaccharide lyases; CE carbohydrate esterases; AA, auxiliary activities; CBM, carbohydrate-binding modules, and SLH, S-layer homology modules. The numeric values in parentheses refer to the numbers of genes of each class. **b**, Number of CAZyme-coding genes in the representative genomes of five rumen ciliate subfamilies and gut fungi. **c**, A PCoA plot showing the CAZyme profiles across the five subfamilies (the color schemes are the same as in Fig 4b), and the P-value of Adonis is < 0.001. **d-e**, The mean number and prevalence of degradative CAZymes encoded in the genomes of rumen bacteria (n = 2,045), gut fungi (n=8), non-gut Alveolate (n =13), and the five subfamilies of rumen ciliates (n =19). **f**, An example for the xylanase (EC 3.2.1.8_GH10) of rumen ciliates originated via horizontal gene transfer (HGT) from rumen bacteria. **g-h**, Structural and activity divergences between rumen ciliates and its bacterial donors for cellulase and xylanase. The symbols *** represent a statistical significance of P-values < 0.001. **i**, Inhibition zone assays of a ciliate lysozyme in GH19.

Predicting the substrates of glycoside hydrolases (GH) and polysaccharide lyases (PL) by categorizing their corresponding EC number (EC 3.2.1.- or EC 4.2.2.-), we found that each species of the rumen ciliates possessed a broad array of enzymes to synergistically degrade plant cell wall (cellulose, hemicellulose and pectin), plant non-structural carbohydrates (fructan and starch) and microbial carbohydrates (chitin and peptidoglycan) (**Fig. 4d**). Moreover, SAGs encoded multiple enzymes to synergistically degrade a single kind of polysaccharides and the resulting oligosaccharides. For example, the Ophryoscolecid SAGs possessed endo-β-1,4-glucanase (EC 3.2.1.4), β-glucosidase (EC 3.2.1.21), endo-1,6-β-glucosidase (EC 3.2.1.75), and β-1,4-cellobiosidase (EC 3.2.1.91), allowing them to degrade cellulose. Less than one-quarter of the degradative CAZymes (ranging from 6% in Ophryoscolecinae to 24% in Isotrichinae) of ciliates were invested in degrading microbial carbohydrates, suggesting that their energy acquisition through microbe predation is limited. Most of the degradative CAZymes of Diplodiniinae (72%) and Ophryoscolecinae (82%) were invested in degrading plant cell wall, while those of Isotrichinae (45%), Dasytrichinae (44%), and Entodiniinae (44%) were mainly invested in degrading plant non-structural carbohydrates (**Fig. 4e**). These observations revealed the molecular mechanisms underlying the variable substrate preferences across subfamilies^6^ and why ciliates could reach up to half of the rumen microbial biomass^6^. The variable investment tradeoffs in degradative CAZymes partly avoid an overlapping niche across subfamilies, and thus enabling a robust performance of the overall biomass degradation process^41^. Additionally, rumen ciliates ingest substrates into digestive vacuoles (**Supplementary Fig. 14**) to which they deliver their degradative enzymes, which can help increase the binding efficiency between degrading enzymes and substrates due to close proximity and improve the utilization efficiency of the degradation products. Taken together, the broad array of CAZymes, variable substrate preferences, and intra-vacuoles digestion in rumen ciliates may contribute to their energy acquisition and use from recalcitrant plant materials and allow for their ecological success in the rumen ecosystem.

Compared with rumen ciliates, non-gut Alveolate species (n =13) possess a much smaller number and lower diversity of degradative CAZymes, and they are incapable of digesting xylan-related hemicellulose (xylan, xyloglucan, and arabinoxylan) or pectin (**Fig. 4d**), suggesting that most degradative CAZymes of rumen ciliates were not likely obtained by vertical inheritance. To determine whether this innovation is the result of inter-kingdom horizontal gene transfer (HGT), we extracted and built phylogenetic trees of homologous sequences from the genomes of rumen bacteria, gut fungi, non-gut Alveolate, and rumen ciliates. The results showed that the CAZymes of rumen ciliates mainly originated via HGT from gut bacteria and fungi (as high amino acid sequence similarity and phylogenetic nesting), and most of them had more than one HGT event, which could occur in the early or during the diversification of rumen ciliates (**Fig. 4f** and **Supplementary Fig. 15**). For example, xylanase (EC 3.2.1.8) has recruited five gene families (GH5, GH10, GH11, GH30 and GH43) from bacterial and fungal donors, involving at least 15 HGT events (9 from Firmicutes, 4 from Bacteroidota, 1 from Fibrobacteria, and 1 from fungi).

Structural divergences of horizontally transferred CAZymes were always observed between ciliates and the corresponding donors, such as the loss of the CBM22 domain in EC 3.2.1.8 in GH43, the acquisition one motif in EC 3.2.1.91 in GH5, the loss of motif in EC 3.2.1.4 in GH5 and EC 3.2.1.8 in GH10, and truncation at terminal non-domain regions of the above four enzymes (**Fig. 4g** and **Supplementary Fig. 16**). To investigate the enzyme activities differences between horizontally transferred CAZymes, we cloned and overexpressed one cellulase (EC 3.2.1.4 in GH5) and one xylanase (EC 3.2.1.8 in GH10) from both bacteria and ciliates using *Pichia pastoris*. The results showed that the ciliate cellulase and xylanase had nine-fold and two-fold higher activity than those of bacterial donors, respectively (**Fig. 4h**).

Additionally, 25.6% of the degradative CAZymes could not be annotated to any EC numbers. We evaluated the substrates and activity of one GH19 protein of *Oph. caudatus* also after cloning and overexpression in *P. pastoris*, and the results showed that it was a lysozyme (with a specific activity of 36,007 U/mg), and the purified GH19 protein inhibited *Micrococcus luteus* (Gram-positive), but not *Escherichia coli* (Gram-negative) (**Fig. 4i**). The characteristic selective inhibition of the Gram-positive bacterium suggests that this ciliate lysozyme was a potential alternative to monensin, which is one of the most successful rumen modifiers in improving ruminant feed efficiency but has been banned in some countries because it is an ionophore and appears residues in milk and meat^42^.

### Enabling metagenomic analyses of rumen ciliates

The sequenced genomes can greatly facilitate metagenomic analyses of microbiomes including communities of rumen ciliates. We re-analyzed 902 publicly available rumen metagenomes (**Supplementary Table 3**) against three datasets: 1) the RefSeq genomes and rumen microbial genomes^2,43^, 2) dataset 1) plus the SAGs of the rumen ciliates, and 3) dataset 2) plus the metagenome-assembled ciliate sequences. The results showed that the read classification rates of metagenome as rumen ciliates could be affected by the host, diet, and management (**Fig 5a, c**). Of note, the classification rates as rumen ciliates decreased by 99% in the metagenomes of sheep that suffered from subacute ruminal acidosis due to feeding of a high concentrate diet^44^, which is consistent with previous experimental findings that ciliates are sensitive to pH^45^. Of the 902 metagenomes, 726 were from animals fed a diet with a concentrate content of <80%, which is considered a low risk for subacute ruminal acidosis according to the dramatic drop of ciliate abundance. And in the 726 rumen metagenomes, our ciliate dataset allowed for classification of 12% of the total reads on average, with a large range from 0.1% to 72% (**Fig 5a**). With the improvement of ciliate dataset, the total read classification rates across metagenomes increased and became more convergent than otherwise. For example, with the improvement of ciliate dataset, the total read classification rates in the 285 samples of Stewart et al.^2,38^ increased from 39-92% to 72-94%, and one-fold more of the samples (33% vs. 73%) achieved a classification rate of 80% or higher (**Fig 5b**). Of the identified ciliate reads in the metagenomes, more than 73% could be classified based on SAGs, suggesting that the collected ciliates comprised a considerable fraction of the rumen ciliates community (**Fig 5d**). Interestingly, based on microscopic counting, Entodiniinae as the most dominant rumen ciliates can account for >90% of the total rumen ciliates in domesticated ruminants (**Supplementary Table 2**), but only ∼25% of the identified ciliate sequence reads were assigned to Entodiniinae. This discrepancy might be attributable to (1) the lack of the genomes of some abundant Entodiniinae species, such as *Ent. dilobum* and *Ent. rostratum*, and (2) the ploidy of Entodiniinae was likely lower than that of other rumen ciliates.

**Fig. 5.**
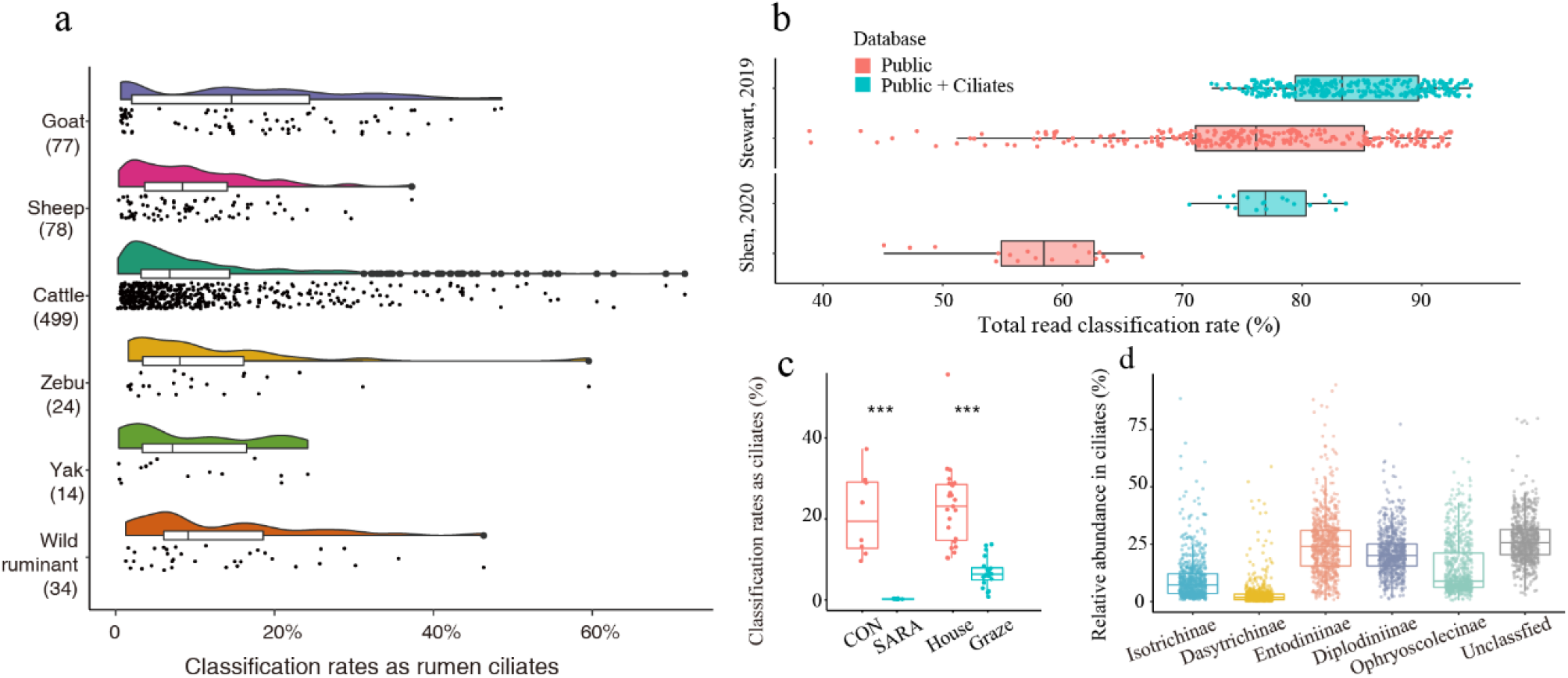
Metagenomic analyses. **a**, The read classification rates as ciliates across rumen metagenomes of different hosts fed a diet with <80% concentrate; **b**, Improvement of classification rates of total reads in two previous studies^2,76^ enabled by the ciliate dataset; **c**, The association of read classification rates as rumen ciliates with diet and management^44,77^. Statistical significance was determined by the two-side Wilcoxon rank-sum test. Sheep were fed diets with low-level concentrate (CON, control, n=8) or high-level concentrate that induced subacute ruminal acidosis (SARA, n=8)^44^. Holstein dairy cows were house fed a corn silage and concentrate diet (n=23) or grazed on a natural prairie (n=20)^77^. **d**, Relative abundance of rumen ciliates.

## Discussion

Compared with gut bacteria and archaea^1–3^, eukaryotic microbes (protozoa and fungi) are understudies and represent an underexplored genetic resource^46^. Here, we used single-cell sequencing to circumvent the lack of the inability to establish axenic cultures of rumen ciliates and developed a ciliate-telomere-capped sequence dataset to aid the identification of ciliate sequences from mixed sequences. Compared with previous studies on fungi and flagellate^47,48^, the more successfully captured ciliate genomes from individual cells were largely attributed to their mini-/nano-chromosomes, which typically have a high ploidy^20,24^. Metagenomic binning is the other technique to recover uncultured microbial genomes^36^, but it is not applicable to recover ciliate genomes due to their large number and nonuniform copy numbers of chromosomes and the dependence of binning on sequencing depth. In this study, the unparalleled genome catalog of rumen ciliates shines new lights on their taxonomy, evolution, and metabolism, and also greatly facilitates the rumen metagenomic analyses by allowing ∼12% of reads to be classified.

How to define the boundaries for individual taxonomic rank for microorganisms including ciliates is challenging^25,26^ because the reproductive isolation principles were not or mostly not applicable and pure cultures are not available for most microorganisms. Combining two quantitative operational methods: ANI and RED, we revised the taxonomy for rumen ciliates, which allowed 45% of the 22 collected morphospecies to be changed to their existing taxonomy. Using this revised taxonomy, nine synonymic Ophryoscolecidae morphospecies, two cryptic Isotrichidae species and two new genera were identified, and genus Dasytricha was reassigned from Isotrichidae to a new family Dasytrichidae. The genome-based taxonomy corroborated or rejected the taxonomic worth of some morphological features of rumen ciliates, which will help to revise the morphology-based taxonomy and to identify new species.

Additionally, we provided a phylogenetic backbone to understand the evolutionary processes of specific traits and early diversification in rumen ciliates, such as the origin of the non-extensively fragmented minichromosomes in Isotrichidae, the origin and loss of skeletal plates in Ophryoscolecidae. The identified novel traits will facilitate future research on ciliates in general. For example, compared with the previous inter-class investigations^49,50^, the rumen ciliates will provide a more allied model for investigating the unsolved molecular mechanisms underlying chromosome fragmentation^21^.

Importantly, we showed that rumen ciliates were another important weapon for ruminants to degrade nearly all kinds of plant-derived polysaccharides. In particular, the species of Diplodiniinae and Ophryoscolecinae encoded as many CAZyme genes as gut fungi, and ∼80% of their degrading enzymes acted on plant cell wall. Most of the ciliate degradative CAZymes were obtained via HGT from bacteria and fungi^7,17^, but they underwent considerable sequence divergence from the donor. The experimentally identified promising activity of their cellulase, xylanase and lysozyme reflects the potential application of ciliate CAZymes in biomass conversion and microbiota modulation.

## Methods

### Single-ciliate-cell isolation, identification and sequencing

All the experimental procedures on animals were conducted per the guidelines of the Regulations for the Administration of Affairs Concerning Experimental Animals (Ministry of Science and Technology, China, 2013) and were approved by the Northwest A&F University Animal Care and Use Committee.

Individual ciliate cells were isolated from fresh rumen fluid samples collected from 15 ruminally fistulated ruminants (6 Holstein cattle, 3 Qinchuan beef steers, and 6 Guanzhong dairy goats) all kept in Yangling, Shannxi Province, China using glass micropipettes under an inverted microscope (Nikon, TI-FL, Japan). We purposely selected ciliates with distinct morphologies. The single ciliate cells were repeatedly transferred and washed using MB9 buffer^51^ until no other ciliate cells could be found microscopically. The washed single ciliate cells were then transferred to individual microtubes each containing 2 µl of cell lysis buffer. A total of 69 single ciliate cells representing 22 morphospecies (as identified morphologically, detailed below) were isolated, and 45 of them were subjected to whole-genome amplification (WGA) using multiple displacement amplification^52^ and the remaining 24 single ciliate cells were subjected to parallel WGA and whole-transcriptome amplification (WTA) using the G&T-seq^22,53^ (**Supplementary Table 1**). Illumina TruSeq libraries were prepared for each of the amplified genomes and transcriptomes and then sequenced (2 ×150 bp) on an Illumina NovaSeq platform.

The morphological identification of rumen ciliates^54^ was conducted with light microscopy, scanning electron microscopy, and confocal laser scanning microscopy. The morphological description and micrographs of the isolated ciliate species are summarized in **Supplementary Fig. 1-3**. The taxonomic identification of the collected single ciliates was confirmed by sequencing analysis of their 18S rRNA genes after the genome assembly. The ciliates in the rumen samples were counted using a Sedgewick-Rafter chamber as described previously^54^.

### Genome assembly and ciliate sequence identification

The cleaned Illumina reads of each WGA sequencing were assembled using both MEGAHIT (v1.2.1)^55^ and SPAdes (v3.13.1)^56^. The contigs assembled by the two assemblers were merged into longer contigs using quickmerge (v1.2.1)^57^.

The sequence characteristics of rumen ciliates were identified as follows. Firstly, the ends (30 bp) of WGA contigs were used to identify telomeric repeats using MEME (v4.12.0)^58^, and the telomeric repeats of the collected morphospecies were all identified as (5’-CCCCAAT)n. Secondly, complete chromosomes (contigs that were capped with at least 1.5 telomeric repeats without mismatch at both ends) were extracted from the WGA contigs using a python script^32^. Contigs with potential contamination sequences (each with a >20% GC content for the subtelomeric region (100 bp immediately adjacent to the telomere) and a GC content >44% GC for the rest of the contig) were filtered out with an error rate of 1% allowed.

The publicly available rumen metagenomes (n=902, **Supplementary Table 3**) were downloaded from the NCBI SRA, and were assembled using MEGAHIT. These contigs were also subjected to decontamination as mentioned above. The decontaminated contigs from the SAGs and the metagenome-assembled contigs were combined into an 18 Gb sequence dataset of rumen ciliate macronuclear sequences, which were capped with telomeric repeats at one or both ends (**Supplementary Fig. 4)**. The 18 Gb of contigs of ciliates plus other rumen microbial genomes^2,36,38–40,43,59,60^ were used as a reference to pick the ciliate sequences that do not have any telomeres in the WGA assembled sequences using blastn (v2.6.0+). The sequences with the best match to ciliate sequences (>85% identity over >500 bp and >50% coverage of reference or query) were considered as ciliate sequences.

The telomereless contigs picked above and the telomere-capped contigs with >3X of read depth were retained in SAGs. Genome completeness of each SAG was assessed using BUSCO v5^61^ with OrthoDB v10^23^ using the predicted protein-coding genes as input. Of the 69 SAGs (**Supplementary Table 4**), 52 had ≥80% of the 171 Alveolata conserved marker genes and were considered “high-quality”, while 17 had <80% of the Alveolata conserved marker genes.

### Gene prediction

The ciliates in rumen fluid samples (n=14) were enriched by filtering (150 µm) and centrifugation (500 ×g for 5 min). The eukaryotic mRNAs in the ciliate-enriched samples were extracted and purified as previously described^61^, and used for metatranscriptome sequencing. The single-cell transcriptomic and metatranscriptomic sequencing reads were mapped to SAGs using Hisat2, and assigned to each species.

Protein-coding genes were *de novo* annotated using AUGUSTUS (v3.2.3)^62^. Firstly, the WTA sequence reads were assembled using Trinity (v2.8.4)^63^. The assembled transcripts were used to train the *de novo* gene prediction model^62^. The best model was that of *Pol. multivesiculatum* with 93.5% sensitivity and 85.7% specificity at the exon level. Gene predictions of SAGs were performed also using AUGUSTUS and the best model with mapped RNA-seq data as ‘‘hints’’. The tRNAs-coding and rRNA genes were identified using tRNAscan-SE (v2.0.5) and Barrnap (v0.9), respectively.

### Circumscribing species

The ANI and alignment coverage between genomes were calculated using pyani (v0.2.10)^64^ with the default parameters. A clear gap existed between 92% and 97% in ANI values, and thus ANI >97% was considered synonym, and ANI <92% was inter-species circumscription.

### Phylogenomic tree construction and higher taxonomic rank normalization

A total of 113 single-copy homologous genes were identified among the 19 representative genomes and *Tet. thermophila* genomes (as outgroup) by OrthoFinder (v2.5.2)^28^. These single-copy genes were aligned and concatenated into supergenes for ML-based phylogenetic using RAxML (v.8.2.9)^65^. A species tree was also constructed using OrthoFinder based on the 52 SAGs.

The concatenated protein-based phylogeny of the 19 rumen ciliates species served as the basis for high taxonomic rank (at and above genus) normalization using the RED values calculated with PhyloRank (v0.1.0)^25^. The RED intervals for normalizing taxonomic ranks were defined as the median RED value for taxa at each rank ±0.1. Comparing the RED values with the RED intervals of taxonomic ranks, the taxa that fell outside of their RED distribution was corrected.

### Divergence time estimation and gene family expansion

Divergence times of these species were estimated based on the ML tree via Bayesian relaxed molecular clock approach using MCMCtree program in the PAML package (v.4.9)^66^. The calibrated points of the common ancestor of rumen ciliate species (<135 Mya) and ophryoscolecid species (<55 Mya) were obtained from Vďačný et al.^5^.

Orthologs gene families among the 52 SAGs were identified using the OrthoFinder. To reduce the limitation of single-cell genomes, the maximum gene number in each gene family among different SAGs of each species was used in the CAFE (v5.0)^67^ to infer the expansion and contraction of gene families.

### Functional annotation

Functional annotation (KEGG, GO and COG) of the predicted proteins was performed using eggNOG mapper (v2.0.1) with the DIAMOND mapping mode, based on the eggNOG 5.0 orthology data^68^. The protein domain was annotated using pfam_scan.pl compared to Pfam_A. CAZymes annotation was executed using dbCAN2^69^ against the CAZyme database V9. The CAZymes were categorized with their corresponding EC number by aligning against uniprot_sprot and CAZyme database V9 using Diamond (v0.9.21.122) with an *E*-value cutoff of e-3.

### Identification of CAZymes with HGT signature

To compare the CAZyme numbers and profiles of rumen ciliates with rumen bacteria (n = 2,405), gut fungi (n=8), non-gut Alveolate (n =13) (**Supplementary Table 5**), the CAZymes of the latter microbes were also annotated as above. The 2,405 rumen bacterial genomes served as the representative genome at species-level (ANI>95%) for the 4,941 metagenome-assembled genomes from Stewart et al.^2^ and 410 genomes from the Hungate 1000 project^1^.

For each CAZyme (the same family with the same EC number), a phylogenetic tree among rumen ciliates and other microbes was constructed using IQ-TREE (v2.1.4-beta)^70^ with 100 bootstrap replicates. The potential HGT events were inferred using the pipeline and cut-offs of Haitjema et al.^39^.

### Catalytic activity assay of cellulase and xylanase

*In vitro* enzymatic activity assays were used to test the the cellulase (EC 3.2.1.4 in GH5) from rumen ciliate (*Oph. caudatus*) and its HGT donor (*Treponema bryantii* NK4A124), and xylanase (EC 3.2.1.8 in GH10) from rumen ciliate (*Ost. gracile*) and its HGT donor (*Ruminococcus flavefaciens* YRD2003). Four genes were codon-optimized according to *P. pastoris* preference. Construction of recombinant plasmids and heterologous expression and purification followed the procedures previously described^71^. The cellulase and xylanase activities were measured as previously described^72,73^. Each experiment was performed with three reaction replicates to determine the mean ± s.d. of the catalytic activity value of the enzymes.

### Catalytic activity assay of lysozyme

The substrates and activity of one GH19 (it includes lysozyme and chitinase) protein of *Oph. caudatus* were examined after cloning into *P. pastoris*. The enzymatic substrate assay showed that it was a lysozyme, which activity were measured as previously described^74^. The antibacterial ability of this lysozyme was tested by inhibition of *Micrococcus luteus* (ATCC 4698) and *Escherichia coli* (CVCC 3367), as a representative of Gram-negative and –positive bacteria, respectively, zone assay on agarose plates.

### Sequencing reads classification

The sequencing reads of 902 publicly available rumen metagenomes were classified using Kraken (v2.0.7-beta)^75^ against three custom databases: 1) Public database, consisting of all microbial genomes from RefSeq (release 201) plus rumen microbial genomes; 2) Public+SAGs database, 3) Public+Ciliates database, consisting of public database, all 52 SAGs, and the ciliate-telomere-capped sequences from the 902 metagenomes.

## Data availability

All sequencing data and assembled genomes have been deposited in the NCBI database with accession ID PRJNA777442. All other relevant data are available in this article and its Supplementary Information files, or from the corresponding author upon request.

## Acknowledgements

This study was financially supported by the National Natural Science Foundation of China (U21A20247, 31822052, to Y.J., and 31902126, to Z.L.) and the China Postdoctoral Science Foundation (2019M663841, to Z.L.). We thank the High-Performance Computing Platform of Northwest A&F University for providing computing resources.

## Author contributions

Z.L. and Y.J. conceived and supervised the project. Z.L., X.W., T.Z., Y.L., T.S. and X.X collected the samples. X.W., Z.L., Y.L., F.L., Y.H., H.H., J.N., and J.T. carried out the experiments. Z.L., Y.Z., X.W., T.Z., X.D., X.P., R.J., Y.Y., and S.G. performed bioinformatic analyses. Z.L. wrote the manuscript. Z.Y., Y.J., H.H., J.Y. and B.Y. revised the manuscript. All authors have read and approved the final manuscript.

## Supplementary figure legend

**Supplementary Fig. 1**. Morphological description and micrographs of the 19 studied rumen ciliate species including the identified synonymic and cryptic species. The micrographs from scanning electron microscopy, light microscopy, and confocal laser scanning microscopy; and the cited schematic diagram78 and morphological description6 are presented. The skeletal plate was stained by Lugo’s iodine. Scale bars, 50 μm (a-x).

**Supplementary Fig. 2**. Summary of rumen ciliates isolated from host animals (**a**) and genome-based subfamilies and genera (**b**).

**Supplementary Fig. 3**. The difference in secondary caudal spines between morphospecies *Oph. caudatus* (three circles) and *Oph. bicinctus* (two circles).

**Supplementary Fig. 4**. A pipeline for recovering single-ciliate genomes and gene prediction. The original array consists of three parts: public metagenomic data, metatranscriptomic data, and single-ciliate genomic and transcriptomic data from whole-genome and whole-transcriptome amplification sequencing.

**Supplementary Fig. 5**. Comparison of genomes characteristics by different sequencing depth and assembly strategies. **a-d**, Contig N50, genome size, the ratio of telomere-capped contigs using different sequencing data size when the genomes were assembled by Megahit. **e-f**. Contig N50 and genome size, the ratio of telomere-capped contigs when the genomes were assembled by different assembly strategies: single software (Megahit, Spades) or hybrid assembly (Cd-hit and Quickmerge). Average size ratio of telomere-capped contigs in Isotrichidae (**g**) and Ophryoscolecidae genomes (**h**) with different assembly strategies.

**Supplementary Fig. 6. a** Pairwise average nucleotide identity (ANI) and coverage comparison calculated for the 52 SAGs. **b** Histogram of pairwise ANI values of morphospecies. **c** Pairwise nucleotide identity of 18S rRNA gene across SAGs.

**Supplementary Fig. 7**. Species tree of 52 rumen single-ciliate genomes inferred using OrthoFinder with the coalescent model.

**Supplementary Fig. 8**. Divergence time estimates within rumen ciliates inferred by MCMCtree. The estimated divergence times with 95% credibility intervals are labeled on each node.

**Supplementary Fig. 9**. Key features of protein-coding chromosomes of Isotrichidae and Ophryoscolecidae, using *Isotricha sp. YL-2021b* and *Polyplastron multivesiculatum* as examples, respectively. Representative chromosome features are shown on average length. UTR, untranslated region; UTS, untranscribed region.

**Supplementary Fig. 10**. Genome characteristics of rumen ciliates. Gene intron length (**a)** and genomic GC content (**b**), gene family richness (**c**) and profiles (**d**) across five subfamilies. **e** Number of gene families along with an additional SAG. **f** Comparison of the gene repertoires of 19 ciliate species. **g** The shared and unique gene families in the five subfamilies of rumen ciliates.

**Supplementary Fig. 11**. The relative chromosome copy number distribution in rumen ciliate genomes.

**Supplementary Fig. 12** Comparison of the KEGG pathway repertoires of rumen ciliates. Significant differences among subfamilies were assessed with Fisher’s exact test. ^a-d^Means with different superscripts within a row differ (P< 0.05).

**Supplementary Fig. 13** Comparison of the Gene Ontology repertoires of rumen ciliates. Significant differences among subfamilies were assessed with Fisher’s exact test. ^a-d^Means with different superscripts within a row differ (P< 0.05).

**Supplementary Fig. 14** Photomicrograph of rumen ciliates were engulfing a plant fiber

**Supplementary Fig. 15** Examples for the CAZymes of rumen ciliates originated via horizontal gene transfer from gut bacteria and fungi or via vertical inheritance from non-gut Alveolate.

**Supplementary Fig. 16** Examples for structural divergences of horizontally transferred CAZymes between ciliates and the corresponding donors

**Supplementary Table 1** Background information on the single-ciliate-cell samples.

**Supplementary Table 2** Relative abundances of rumen ciliates across our collected samples and published studies as evaluated by microscopic counting.

**Supplementary Table 3** Characteristics and sources of rumen metagenomic datasets used in this study.

**Supplementary Table 4** The statistic information of the single-ciliate-cell amplified genomes.

**Supplementary Table 4** The statistic information of the single-ciliate-cell amplified genomes.

**Supplementary Table 5** The genomes of gut fungi (n=8) and non-gut Alveolate (n=13) used in this study

